# Ecology and age, but not genetic ancestry, predict fetal loss in a wild baboon hybrid zone

**DOI:** 10.1101/2022.09.03.505836

**Authors:** Arielle S. Fogel, Peter O. Oduor, Albert W. Nyongesa, Charles N. Kimwele, Susan C. Alberts, Elizabeth A. Archie, Jenny Tung

**Author notes:** Corresponding author Correspondence: Jenny Tung, Department of Primate Behavior and Evolution, Max Planck Institute for Evolutionary Anthropology, Deutscher Platz 6, 04103 Leipzig, Saxony, Germany. Contributed equally to this work (Arielle S. Fogel, Peter O. Oduor).

## Abstract

**Objectives:** Pregnancy failure and fetal loss represent a major fitness cost for any mammal, particularly those with slow life histories such as primates. Here, we quantified the risk of fetal loss in wild hybrid baboons, including genetic, ecological, and demographic sources of variance. We were particularly interested in testing the hypothesis that hybridization imposes a cost by increasing fetal loss rates. Such an effect would help explain how baboons maintain taxonomic integrity despite interspecific gene flow.

**Materials and Methods:** We analyzed pregnancy outcomes for 1,020 pregnancies observed over 46 years in a natural yellow baboon-anubis baboon hybrid zone. Fetal losses and live births were scored based on near-daily records of female reproductive state and the appearance of live neonates. We modeled the probability of fetal loss as a function of a female’s genetic ancestry (based on whole-genome resequencing data), age, number of previous fetal losses, dominance rank, group size, climate, and habitat quality using binomial mixed effects models.

**Results:** Female genetic ancestry did not predict the likelihood of fetal loss. Instead, the risk of fetal loss is elevated for very young and very old females. Fetal loss is most robustly predicted by ecological factors, including poor habitat quality and extreme heat during pregnancy.

**Discussion:** Our results suggest that gene flow between yellow baboons and anubis baboons is not impeded by an increased risk of fetal loss for hybrid females. Instead, ecological conditions and female age are key determinants of this component of female reproductive success.

**Research Highlights:**

- Female baboons do not experience fetal loss as a cost of hybridization.
- Heat stress, poor habitat quality, and young and old age elevate the risk of fetal loss, emphasizing roles for ecology and life history in determining birth outcomes.

**Graphical Abstract:** Neonate drawings by Emily Nonnamaker.

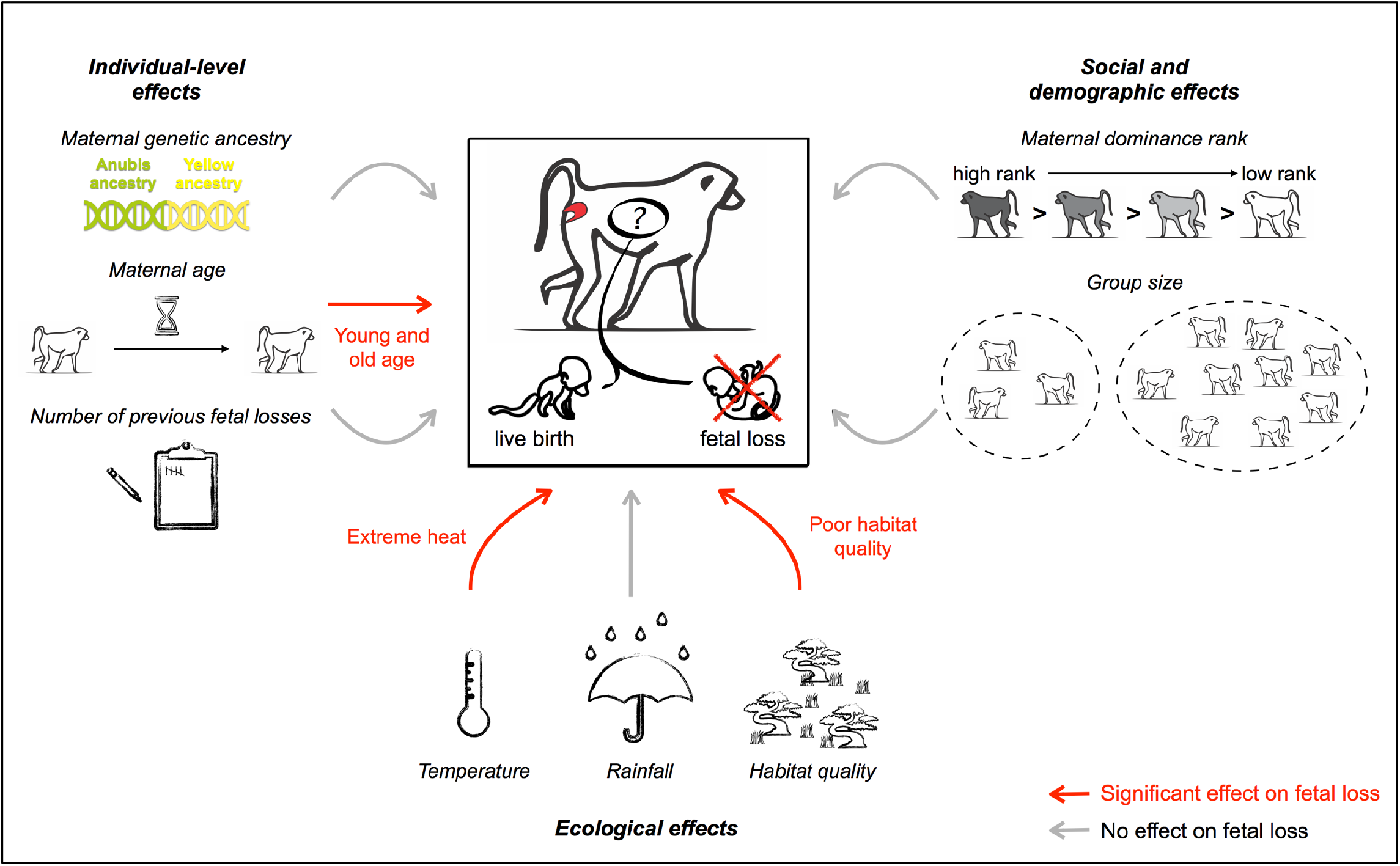

## Main Text

Hybridization (i.e., interbreeding between distinct genetic lineages) is a common feature of primate evolution. Historic or ongoing hybridization has been documented in all primate families, including in the genus *Homo* (reviewed in Arnold & Meyer, 2006; Dannemann & Racimo, 2018; Tung & Barreiro, 2017; Zinner, Arnold, & Roos, 2011). In many cases, however, hybridizing taxa remain phenotypically and genetically distinct, and the costs of hybridization are often subtle or have not yet been detected. These observations defy the expectation that hybridization should have a homogenizing effect and ultimately lead to the erosion of species differences. They also contrast with cases of reduced hybrid fertility and/or viability reported in many non-primate hybrids (e.g., Ålund, Immler, Rice, & Qvarnström, 2013; Neubauer, Nowicki, & Zagalska-Neubauer, 2014; Powell et al., 2020; Svedin, Wiley, Veen, Gustafsson, & Qvarnström, 2008; Turner, Schwahn, & Harr, 2012; Walsh et al., 2016). Thus, the mechanisms that limit gene flow between primate lineages remain a major unresolved puzzle in the study of primate biodiversity and evolution.

In taxa that interbreed yet maintain their taxonomic distinctiveness, hybrid offspring are predicted to incur fitness costs. Costs to fertility are a likely candidate, given the apparent viability of many natural hybrids and the potential subtlety of effects on fecundability or miscarriage rates. Indeed, in captivity, crosses among *Eulemur*, tamarin, and macaque species produce hybrid offspring with compromised fertility (Bernstein & Gordon, 1980; Rumpler & Dutrillaux, 1980; Soto-Calderón et al., 2018; Tattersall, 1993). A few studies of naturally occurring admixture also provide preliminary support for fertility-related isolating barriers. For example, in a hybrid zone in Mexico between two distantly related howler monkey species, crosses between mantled howler females (*Alouatta palliata*) and black howler males (*A. pigra*) do not appear to produce fertile offspring (Cortés-Ortiz et al., 2007). In more recently diverged chacma baboons (*Papio ursinus*) and Kinda baboons (*P. kindae*) in Zambia, the rarity of offspring from crosses between small female Kindas and large male chacmas may be a result of gestational and obstetric challenges that limit gene flow, at least in one direction (Jolly, Burrell, Phillips-Conroy, Bergey, & Rogers, 2011). Finally, in the human lineage, reduced fertility has been hypothesized to have limited admixture between the ancestors of modern humans and Neanderthals and Denisovans (Jégou, Sankararaman, Rolland, Reich, & Chalmel, 2017; Sankararaman et al., 2014; Sankararaman, Mallick, Patterson, & Reich, 2016). However, in all three of these cases, evidence for compromised fertility in hybrids is indirect because no phenotypic data on fertility-related traits is available.

Baboons, members of the genus *Papio*, are well-suited for assessing fertility-related costs to hybrids as they are intensively studied in the wild, have external indicators of reproductive state that facilitate data collection on mating and pregnancy outcomes, and frequently hybridize in nature (Altmann, 1973; Fischer et al., 2019). Originating in southern Africa, baboons subsequently expanded across the continent 1-2 million years ago to form two distinct lineages: the northern clade, including the anubis (or olive) baboon (*P. anubis*), the hamadryas baboon (*P. hamadryas*), and the Guinea baboon (*P. papio*); and the southern clade, including the chacma baboon (*P. ursinus*), the Kinda baboon (*P. kindae*), and the yellow baboon (*P. cynocephalus*) (Rogers et al., 2019). Today, these species occupy largely non-overlapping geographic ranges across Africa and the Arabian peninsula and are distinguishable based on morphological and behavioral features (Fischer et al., 2019). However, although current scientific consensus recognizes them as distinct species (Fischer et al., 2019; Rogers et al., 2019), they interbreed to produce hybrids at the boundaries between their current geographic ranges (Charpentier et al., 2012; Jolly et al., 2011; Phillips-Conroy & Jolly, 1986). This process of interspecific exchange appears to be part of a long, complex history of independent evolution interspersed with repeated episodes of gene flow among baboon species (Rogers et al., 2019; Vilgalys et al., 2022; Wall et al., 2016; Zinner, Groeneveld, Keller, & Roos, 2009).

What keeps baboon species distinct? Studies of natural hybrid zones—between anubis and hamadryas baboons in Ethiopia, chacma and Kinda baboons in Zambia, and yellow and anubis baboons in Kenya—have not yet produced clear answers. In Ethiopia, anubis baboons hybridize with hamadryas baboons despite major differences in their social and mating structures: multilevel societies featuring one-male, multifemale units and female dispersal in hamadryas baboons, versus polygynandrous, multimale, multifemale social groups with male dispersal in anubis baboons (Bergman & Beehner, 2004; Kummer, 1968; Kummer, Götz, & Angst, 1970). These species differences may impose costs on hybrids, and in support, early reports suggested that hybrid males employ ineffective mating strategies (Nagel, 1970). However, more recent work indicates that hybrid males may enjoy high reproductive success in highly admixed social groups (Bergman, Phillips-Conroy, & Jolly, 2008). Meanwhile, in the Zambian hybrid zone between Kinda baboons and chacma baboons, asymmetric hybridization also suggests potential costs of hybridization (Jolly et al., 2011). Both morphological and behavioral mechanisms have been hypothesized, but their roles in mediating reproductive isolation-associated costs have not been quantified.

In the well-characterized baboon hybrid zone between yellow baboons and anubis baboons in East Africa, hybrids are viable and reproduce readily with both parent species and with other hybrids, even though their parent species are as distantly related as possible among extant baboons (∼1.4 million years diverged vs. ∼750 thousand years for anubis-hamadryas and ∼600 thousand years for chacma-Kinda) (Rogers et al., 2019). Yellow and anubis baboons have similar social systems and are similar in size, so neither fundamental differences in mating behavior nor gestational and obstetric considerations likely pose a barrier to hybridization (Fischer et al., 2019; Rogers et al., 2019). Indeed, yellow-anubis hybrids may sometimes experience phenotypic advantages. In the Amboseli region of Kenya, near the center of the hybrid zone, greater anubis ancestry in this majority yellow ancestry population is associated with accelerated maturation and an increased rate of opposite sex affiliation, an important predictor of lifespan in this population (Archie, Tung, Clark, Altmann, & Alberts, 2014; Campos, Villavicencio, Archie, Colchero, & Alberts, 2020; Charpentier, Tung, Altmann, & Alberts, 2008; Fogel et al., 2021). In males, more anubis ancestry also predicts earlier natal dispersal and increased mating success (Charpentier et al., 2008; Tung, Charpentier, Mukherjee, Altmann, & Alberts, 2012). Nevertheless, the hybrid zone is narrow relative to the large geographic ranges of the two parent species (Charpentier et al., 2012) and genomic analyses reveal selection against admixture in Amboseli (Vilgalys et al., 2022). Together with evidence that gene flow between yellow baboons and anubis baboons is a repeated occurrence in baboon evolutionary history (Rogers et al., 2019; Vilgalys et al., 2022; Wall et al., 2016), these observations suggest that fitness costs to admixture must exist, but are likely subtle and/or temporally or spatially variable.

Here, we investigated whether fertility-related costs, measured in terms of fetal loss (miscarriage or stillbirth), act as a mechanism to restrict interspecific gene flow in the Amboseli baboon hybrid zone. To do so, we combined the most comprehensive data set yet compiled for cases of fetal loss in wild non-human primates (n=1,020 pregnancies in 175 baboon females) with recent estimates of genetic ancestry derived from whole-genome resequencing data (Vilgalys et al., 2022). We also placed these genetic ancestry effects in the context of other variables known or suspected to contribute to fetal loss in primates, including age, dominance rank, group size, ecological conditions (e.g., rainfall, temperature, and overall habitat quality), and individual history of miscarriage (e.g., Bailey, Eberly, & Packer, 2021; Beehner, Onderdonk, Alberts, & Altmann, 2006; Dezeure, Charpentier, & Huchard, 2022; Kolte, Westergaard, Lidegaard, Brunak, & Nielsen, 2021; Nybo Andersen, Wohlfahrt, Christens, Olsen, & Melbye, 2000; Packer, Collins, Sindimwo, & Goodall, 1995; Robbins, Robbins, Gerald-Steklis, & Steklis, 2006; Roof, Hopkins, Izard, Hook, & Schapiro, 2005; Schlabritz-Loutsevitch et al., 2008; Wasser, 1995).

## Materials and methods

### Study site and subjects

The Amboseli basin of southern Kenya is a semi-arid, savanna environment situated within a dry Pleistocene lakebed near the northern base of Mt. Kilimanjaro. Animals in this ecosystem contend with “a place of extremes” (Alberts, 2019). Temperatures vary from extreme midday heat, which can reach 45ºC during the hottest months of the year, to nighttime lows of 5ºC during the coldest months. Rainfall is generally absent during the predictable long dry season from June to October and is extremely variable and unpredictable during the long wet season from November to May. On average, annual rainfall totals equal approximately 350 mm, but rainfall is also highly variable across years (Alberts, 2019)

Subjects in this study were adult females from multiple social groups of wild baboons living in the Amboseli ecosystem. This baboon population has been intensively studied for over five decades, revealing both substantial variation in pregnancy outcomes and a complex history of admixture (Alberts & Altmann, 2001; Alberts & Altmann, 2012; Beehner, Onderdonk, et al., 2006; Samuels & Altmann, 1986; Tung, Charpentier, Garfield, Altmann, & Alberts, 2008; Vilgalys et al., 2022; Wall et al., 2016). All animals in this majority-yellow baboon population are multigenerational hybrids: some individuals harbor anubis ancestry from hybridization events that predate long-term observations (hereafter, historic hybrids), while others are products of both historic gene flow and a recent wave of admixture dating from the 1980’s (hereafter, recent hybrids) (Samuels & Altmann, 1986; Tung et al., 2008; Vilgalys et al., 2022; Wall et al., 2016). The Amboseli hybrid population is located close to the center of a narrow yellow-anubis hybrid zone in southern Kenya that minimally extends into central Kenya and likely occurs wherever anubis and yellow baboon ranges meet (Charpentier et al., 2012; Maples & McKern, 1967).

Members of the study population are individually recognizable and followed on a near-daily basis. During these follows, data are collected on individual-level behavior and reproductive status as well as group demography (Alberts, Archie, Altmann, & Tung, 2020). Study subjects for this analysis were adult females followed between November 1976 and December 2021 as members of 23 different social groups, representing two original study groups and their subsequent fission and fusion products. A minority of these groups (3 out of 23) were semi-provisioned because of their close proximity to a tourist lodge. While these three groups differ demographically and behaviorally from wild-feeding groups (e.g., they exhibit reduced male dispersal, higher rates of inbreeding, and shorter inter-birth intervals: Altmann & Alberts, 2003; Galezo et al., 2022), our model results were similar whether we included or excluded them. Therefore, we included pregnancies from all social groups in our main analysis and we report results that exclude subjects in food-supplemented groups in the Supplementary Information (Table S1).

We restricted the data set to females for whom genetic ancestry estimates from whole-genome resequencing data were available (Vilgalys et al., 2022) and whose birthdates were known within ± 6 months’ error. We also excluded one female who was a known reproductive outlier in the study population (i.e., she experienced continuous cycling and failed to conceive over many years). The resulting sample contained 175 unique females, who together were observed during 1,020 pregnancies (see below).

The research in this study was approved by the Institutional Animal Care and Use Committee (IACUC) at Duke University (#A273-17-12). In Kenya, our research was approved by the Kenya Wildlife Service (KWS), the National Environment Management Authority (NEMA), and the National Council for Science, Technology, and Innovation (NACOSTI).

### Fetal losses

To track pregnancy outcomes, we relied on an established method regularly used during long-term monitoring of this population (Alberts et al., 2020; Altmann, 1973; Beehner, Nguyen, Wango, Alberts, & Altmann, 2006). Briefly, pregnancy is detectable when a female (i) ceases sexual cycling (i.e., ceases to exhibit sex skin swellings and does not menstruate) and (ii) her paracallosal skin gradually changes from black to pinkish red (the “pregnancy sign” in baboons: Altmann, 1973). A conception date is then estimated *a posteriori* as the first day of deturgescence of her sexual swelling during the cycle in which conception occurred (i.e., the conceptive cycle). Analyses of steroid hormone profiles from reproductive females in our study population confirm that this visual assessment method identifies 97% of endocrinologically-identified pregnancies (Beehner, Nguyen, et al., 2006). However, we are likely to miss pregnancies that terminate early in gestation (e.g., in the first trimester, and especially early in the first trimester).

A pregnancy ends with either a live birth—when a previously pregnant female is seen with a new infant—or a fetal loss. Fetal losses are recorded when a female who has been scored as pregnant on the basis of the above criteria resumes cycling without producing a live infant, and also shows signs of fetal loss that may include vaginal bleeding, production of a dead fetus, and/or hormonal signatures of fetal loss (see Beehner, Nguyen, et al. (2006) for a detailed description and validation of these methods).

We excluded pregnancies that ended due to the pregnant female’s death. We included all pregnancies that overlapped periods when social groups were fissioning or fusing, as well as all pregnancies that occurred during several periods of reduced data collection. We also included pregnancies that occurred during the 2009 hydrological year (November 1^st^, 2008 to October 31^st^, 2009), which included the most severe drought ever recorded in the Amboseli ecosystem and that led to reduced conception rates (Carabine, Wainwright, & Twyman, 2014; Lea, Altmann, Alberts, & Tung, 2015; Okello et al., 2016; Tuqa et al., 2014). Models that excluded these time periods produced similar results, however, as reported in Table S1.

### Genetic ancestry

Genome-wide estimates of admixture—here, the estimated proportion of a study subject’s genome derived from anubis ancestry—were included for all females. These estimates can range from 0 (unadmixed yellow) to 1 (unadmixed anubis) and were based on low coverage whole-genome resequencing data generated in a previous study (median = 1.08x coverage, mean = 2.00x coverage) (Vilgalys et al., 2022). To estimate genome-wide ancestry, we first generated local ancestry calls across the genome using a composite likelihood method suitable for low coverage data (*LCLAE*: Wall et al., 2016) and parental species’ allele frequencies for putatively unadmixed yellow and anubis baboons (Robinson et al., 2019; see Vilgalys et al., 2022). We then averaged local ancestry calls across the autosomes for each female to produce a global, genome-wide estimate. Females in this study varied in their ancestry estimates from 0.23 to 0.60 (mean ± SD: 0.36 ± 0.08; Fig. 1).

**Figure 1.**
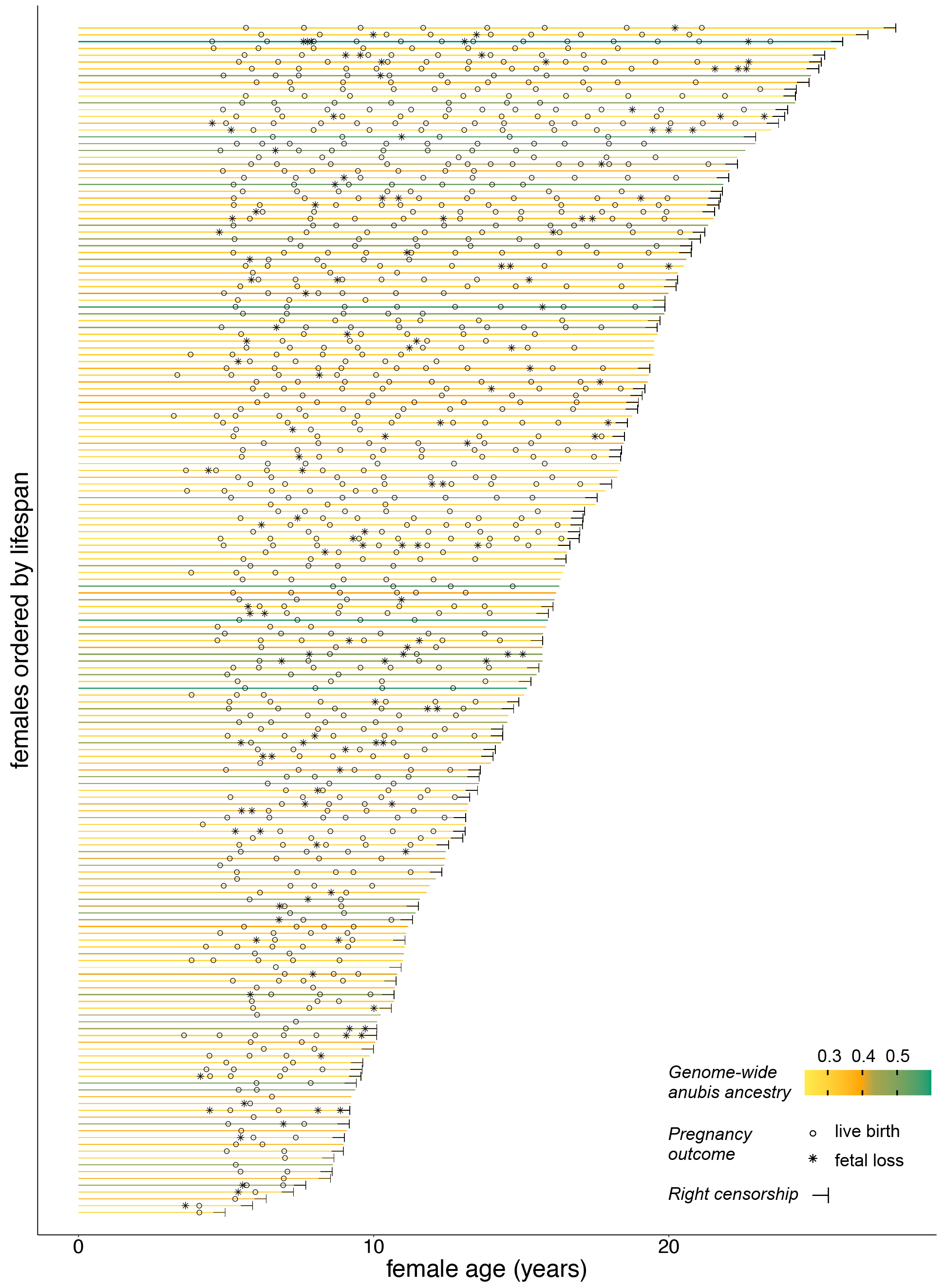
Live birth and fetal loss patterns in Amboseli baboon females. Each horizontal line represents the observed lifespan of a single female. The right end of each line corresponds to either the female’s age at last observation (right censorship; denoted by a horizontal T) or age at death. Horizontal lines are organized from bottom to top based on lifespan and are colored by genetic ancestry. Open circles show live births and asterisks show fetal losses.

To assess genetic ancestry effects on fetal loss, we included both linear and quadratic effects of genome-wide ancestry in our model. The linear effect tests the hypothesis that females with more anubis ancestry in this majority yellow population have an increased likelihood of fetal loss. The quadratic effect tests the hypothesis that intermediate hybrids are more likely to experience fetal loss than animals similar to either parental species, as might be expected if F1-like animals incur the greatest costs of admixture. We also reasoned that admixture-related costs might not be detectable in historically admixed animals if selection has had sufficient time to remove deleterious variants. Therefore, we ran an alternative model in which we replaced the continuous estimates of ancestry with a binary variable corresponding to whether a female was a recent (n=89) or historic (n=75) hybrid (see Vilgalys et al., 2022). Because we could not assign historic versus recent hybrid status for 11 females in our sample, this model was fit to a reduced data set (164 females, 944 pregnancies, 136 fetal losses).

### Ecological effects

Even in non-seasonal breeders like baboons, ecological conditions can still influence female reproductive timing and pregnancy outcomes (Altmann & Alberts, 2003; Beehner, Onderdonk, et al., 2006; Bercovitch & Harding, 1993; Gesquiere, Altmann, Archie, & Alberts, 2018; Hill, Lycett, & Dunbar, 2000; Lea et al., 2015; Lycett, Weingrill, & Henzi, 1999). We therefore evaluated the effects of three key aspects of baboon ecology on the probability of fetal loss: temperature, rainfall, and habitat quality.

### Temperature

Heat stress, when the environment drives core body temperature above its optimum, can have deleterious effects on sperm production, oocyte maturation, and fetal and placental growth (reviewed in Boni, 2019; Hansen, 2009; Walsh et al., 2019). To assess the potential effects of heat stress on fetal loss, we followed Beehner, Onderdonk, et al. (2006) by assessing the effects of temperature in two time periods: the two months prior to conception and the two months prior to the live birth or fetal loss. Temperature (ºC) was measured daily using a min-max thermometer. We calculated the average daily maximum temperature for both time periods, which were weakly negatively correlated, indicating that females who conceived during the cooler months of the year tended to experience pregnancy outcomes in the warmer months of the year and vice versa (Pearson’s *r* = -0.157, P < 10^−6^; Table S2). This is expected given patterns of seasonality in Amboseli and the 6-month gestation period of baboons.

For daily maximum temperature data collected from June 1992 to 1997, the thermometer was located close to the research camp’s kitchen, producing systematically high temperature measurements. Before and after that time period, the thermometer was placed in several other locations in the research camp that were far from any building structure. To correct the data from June 1992 to 1997, we subtracted 4.2ºC from all maximum daily temperature values recorded during this time period. This adjustment factor was calculated based on modeling daily maximum temperature as a function of the day of the year and a random effect of thermometer (instrument used from 1992-1997 versus the four other thermometers). Importantly, a model excluding pregnancy records from this time period produced qualitatively unchanged results.

### Rainfall

In the semi-arid Amboseli ecosystem, rainfall mediates female fertility by affecting food availability and thus female nutritional condition. Following Beehner, Onderdonk, et al. (2006), we evaluated the effects of rainfall conditions in the five months prior to conception as well as the five months prior to the pregnancy outcome (either live birth or fetal loss). Daily rainfall (millimeters) was measured every morning using a rain gauge. For both the five months preceding conception and the end of the pregnancy, we calculated the mean daily rainfall across each time period. These two predictors were moderately negatively correlated in our data set, again consistent with seasonal rainfall patterns in Amboseli (Pearson’s *r* = -0.385, P < 10^−36^; Table S2).

### Habitat quality

In the central part of the Amboseli basin, the mid-1960s to the mid-1980s saw a precipitous decline in *Acacia* woodlands (Altmann, Hausfater, & Altmann, 1985; Western & Van Praet, 1973). Because baboons in Amboseli rely on *Acacia* trees as an important food source and for sleeping sites, this change led to substantially degraded habitat quality for them (Altmann et al., 1985). Subsequently, in the late 1980s and early 1990s, the two study groups then monitored by the Amboseli Baboon Research Project shifted their home ranges ∼5-6 km from the central part of the basin to its south-western perimeter, where food was more readily available (Altmann & Alberts, 2003; Bronikowski & Altmann, 1996). After the home range shift, the baboons spent more time resting and socializing and less time foraging, and females experienced earlier maturation, increased offspring survival, and shorter interbirth intervals (Alberts et al., 2005; Altmann & Alberts, 2003; Bronikowski & Altmann, 1996; Gesquiere et al., 2018). We therefore modeled habitat quality as a binary variable in our model, indicating whether a pregnancy was conceived pre-or post-home range shift (i.e., in low or high habitat quality, respectively).

### Other potential sources of variance in fetal loss rates

### Age

Female age is a known predictor of fertility in a diverse set of species (e.g., Campos et al., 2022; Ericsson, Wallin, Ball, & Broberg, 2001; Froy, Phillips, Wood, Nussey, & Lewis, 2013; Gruhn et al., 2019; Hayward et al., 2013; Jones et al., 2014; Nussey et al., 2009; Reid, Bignal, Bignal, McCracken, & Monaghan, 2003). In the Amboseli baboons, conception probabilities peak in mid-adulthood, and younger and older females experience the longest periods of cycling before conceiving and the shortest pregnancy lengths (Beehner, Onderdonk, et al., 2006; Campos et al., 2022; Gesquiere et al., 2018). We therefore included linear and quadratic effects of female age at conception in our model (following Beehner, Onderdonk, et al., 2006; Campos et al., 2022; Gesquiere et al., 2018). Because female age was highly correlated with parity (primiparous versus multiparous) (Pearson’s *r* = 0.515, P < 10^−69^), we included female age but not parity in the model. For 92% (161 out of 175) of the females in the data set, birthdates were known to within a few days’ error. For the remaining female subjects, 12 females had birthdates that were estimated to within ± 3 months and 2 females had birthdates that were estimated to within ± 6 months (see Table S1 for results of a model excluding females with birthdates estimated with greater than a few days’ error, which were similar to the main analysis). Female age in our data set ranged from 3.2 to 23.4 years of age, with a mean female age of 10.4 years.

### Social status

In social animals, dominance rank can dictate a female’s access to a variety of crucial resources including food, mates, and social partners, which may then affect her reproductive success (e.g., Holekamp, Smale, & Szykman, 1996; Pusey, Williams, & Goodall, 1997; Setchell, Lee, Wickings, & Dixson, 2002; von Holst et al., 2002; Wasser, Norton, Kleindorfer, & Rhine, 2004; Wright et al., 2020; reviewed in Stockley & Bro-Jørgensen, 2011). We therefore modeled a female’s rank at the time of conception as a fixed effect in the model.

Ordinal ranks were assigned on a monthly basis using observed wins and losses in agonistic interactions between all pairs of adult females living in the same social group in a given month: the top-ranking female in the hierarchy in each month was assigned rank 1 and lower-ranking individuals are assigned successively higher numbers (i.e., ranks 2, 3, 4 … *x*, where *x* is the total number of adult females in the social group) (Alberts et al., 2020; Alberts & Gordon, 2018).

The ordinal ranking approach assumes that rank-based competition for resources is density-dependent, such that the resources over which females compete do not scale with changes in group size (Levy et al., 2020). However, females may also compete for density-independent resources, which is better captured by proportional rank (Levy et al., 2020).

Therefore, we also assessed the robustness of our models to substituting proportional rank for ordinal rank. Proportional rank was calculated using ordinal rank values and hierarchy size, such that values range from 0 to 1 and represent the proportion of females in the hierarchy that a given female outranks (i.e., the highest-ranking female is assigned a value of 1 because she outranks 100% of other individuals in the hierarchy). Using ordinal or proportional ranks as the rank-related predictor in our model did not qualitatively change our results so we report results using ordinal rank in the main text (see Table S3 for results using proportional rank).

### Group size

To measure group size, which indexes experienced density and resource competition, we included the number of adult females in the social group at the time of conception as a continuous predictor in our model. We used the number of adult female group members instead of the total number of group members because the number of adult females in the social group is the stronger predictor of female fertility traits, such as inter-birth intervals, in the Amboseli baboons (Altmann & Alberts, 2003)

### Previous fetal losses

The number of previous fetal losses is one of the two strongest predictors of miscarriage rates in humans (in addition to maternal age) (Brosens et al., 2022). We therefore included the number of previous fetal losses as a fixed effect predictor when modelling pregnancy outcome. Primiparous mothers were assigned a value of 0 (no previous fetal losses).

### Statistical analysis

We analyzed the probability of fetal loss using a mixed effects logistic regression approach. Each row of data corresponded to a unique pregnancy and was assigned a value of 1 if the pregnancy resulted in a fetal loss and a 0 if the pregnancy resulted in a live birth. We fit the model using the R package *glmmTMB* (Brooks et al., 2017):

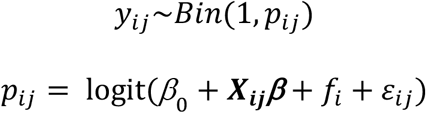

where *y*_*ij*_ is 0 or 1, corresponding to whether female *i* experienced a live birth (0) or fetal loss (1) during pregnancy *j. y*_*ij*_ is drawn from a binomial distribution, where the probability of fetal loss (*p*_*ij*_) is modeled as the function of the logit-transformed sum of (i) the intercept, *β*_*0*_; (ii) the fixed effects (***X***_***ij***_***β***) of female genome-wide ancestry (or recent versus historic hybrid status), female genome-wide ancestry squared, female age at conception, female age at conception squared, number of previous fetal losses, female ordinal dominance rank, group size, average daily maximum temperature two months prior to conception, average daily maximum temperature two months prior to live birth/fetal loss, average daily rainfall five months prior to conception, average daily rainfall five months prior to live birth/fetal loss, and habitat quality (*X*_***ij***_ represents all of these data using standard matrix notation and ***β*** refers to the vector of all fixed effect estimates); and (iii) the random effect of female identity, *f*_*i*_. β_*ij*_ represents model error. All analyses were run and figures were made in R (v.3.6.1; R Core Team, 2019).

### Data and code availability

Data on pregnancy outcomes and all predictor variables tested in our models are available in Table S4. R code for recreating the analyses and figures in the manuscript is available at https://github.com/ArielleF/Fetal-Loss and will be archived on Zenodo upon publication.

## Results

### Fetal loss rates in the Amboseli baboon population

The resulting data set consisted of 1,020 pregnancies (minimum number of pregnancies per female = 1, maximum = 18, mean = 6). For 98% of pregnancies, the dates of conception (1,002 out of 1,020) and end of the pregnancy (1,005 out of 1,020) were known to within a few days’ error. Inclusion or exclusion of pregnancies with less certain conception or pregnancy end dates did not qualitatively change our model results (Table S1). Of the 1,020 pregnancies in our data set, 143 (14%) resulted in fetal loss (see Beehner, Onderdonk, et al. (2006) for similar results in a smaller, earlier dataset for this population). Thus, the rate of fetal losses in our population has been relatively stable over the course of our long-term study, and is similar to the rate reported in human females following clinical recognition of pregnancy (i.e., usually after 4-6 weeks gestational age) (Dimitriadis, Menkhorst, Saito, Kutteh, & Brosens, 2020; Pinar, Gibbins, He, Kostadinov, & Silver, 2018). Exactly half of the females in our sample never experienced a fetal loss, and both the number of fetal losses and number of live births per female tracked their total number of pregnancies (linear model estimate for live births: β = 0.824, p < 10^−89^, Fig. S1A; linear model estimate for fetal losses: β = 0.176, p < 10^−14^, Fig. S1B). Fetal losses were recorded during all three trimesters, although almost half were documented during the third trimester (Fig. 2). Very few fetal losses were related to the immigration of feticidal males (n=4-8 of the 113 fetal losses in our data set that overlapped the data set of Zipple et al. (2017); minimum value corresponds to high confidence feticides and maximum value includes additional possible feticides), so we included these cases in our main data set. A model excluding feticides is consistent with the results from the main model (Table S5).

**Figure 2.**
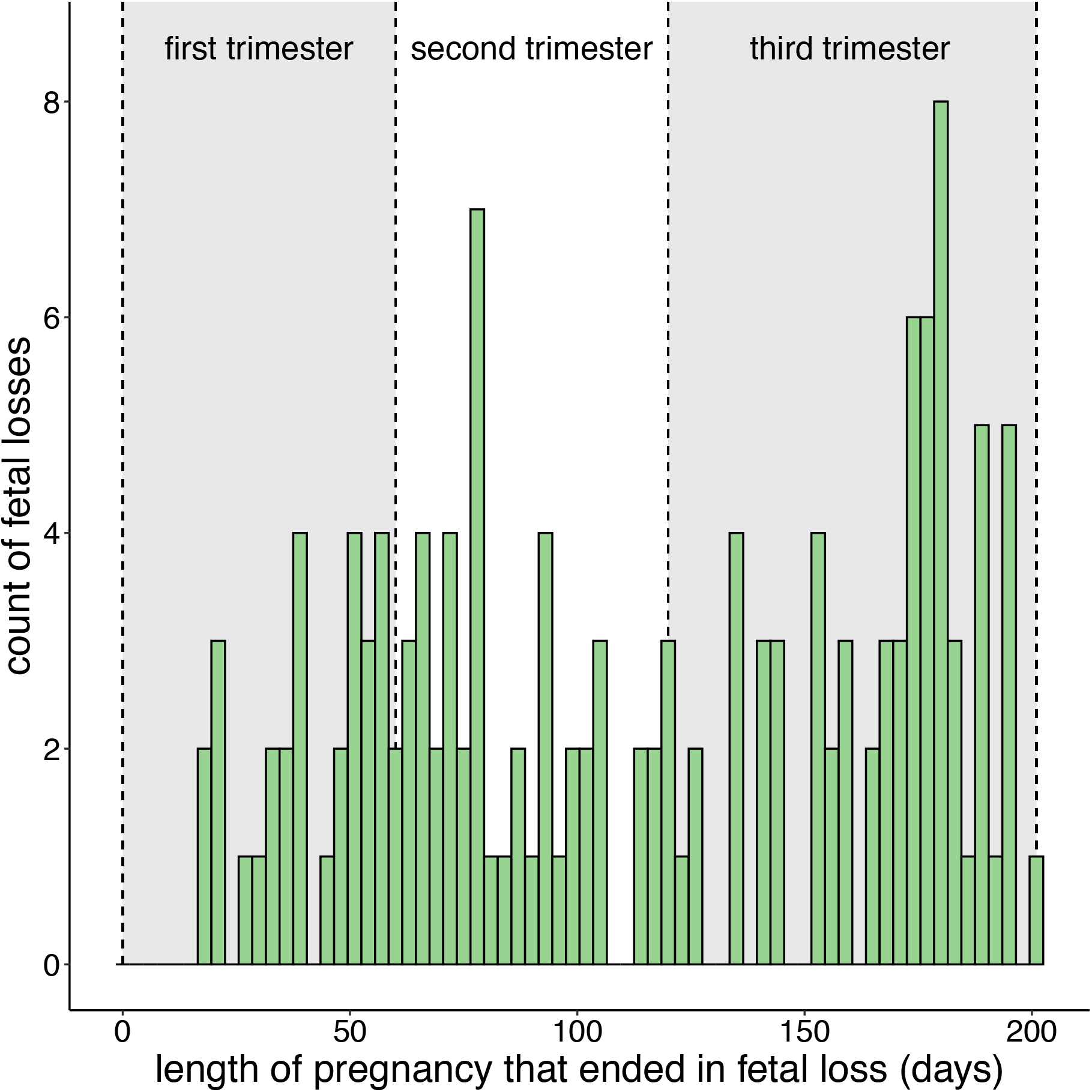
Records of fetal loss by trimester in the data set (n=143). Approximately half of observed fetal losses (67 out of 143) occurred in the third trimester. Mean gestation length for live births in baboons is 178 days (Gesquiere et al., 2018), but stillbirths sometimes occurred several weeks beyond this mean value; thus, the third trimester losses in this dataset occurred from days 121 – 202 post-conception (where 202 is the maximum number of days at which fetal loss was documented). Second and first trimester losses represented 32.9% (47 out of 143) and 20.3% (29 out of 143) of fetal losses, respectively. Data are binned in three-day increments and include 3 cases where the dates of conception or end of the pregnancy were known to greater than a few days’ error.

### Genetic ancestry and hybrid status are not associated with fetal loss

The results of our main model indicate that a female’s genome-wide ancestry did not predict her likelihood of fetal loss (linear effect: β = -8.492, P = 0.506; quadratic effect: β = 11.454, P = 0.471; Table 1, Fig. 1, Fig. 3A). Substituting a binary variable corresponding to a female’s hybrid status (i.e., recent or historic) in place of the linear and quadratic genetic ancestry predictors also did not reveal any effect of admixture on pregnancy outcome (β = 0.201, P = 0.408; Table S6). Both models produced qualitatively similar results and while we focus on the main model below, we note when results differed between models.

**Table 1.**
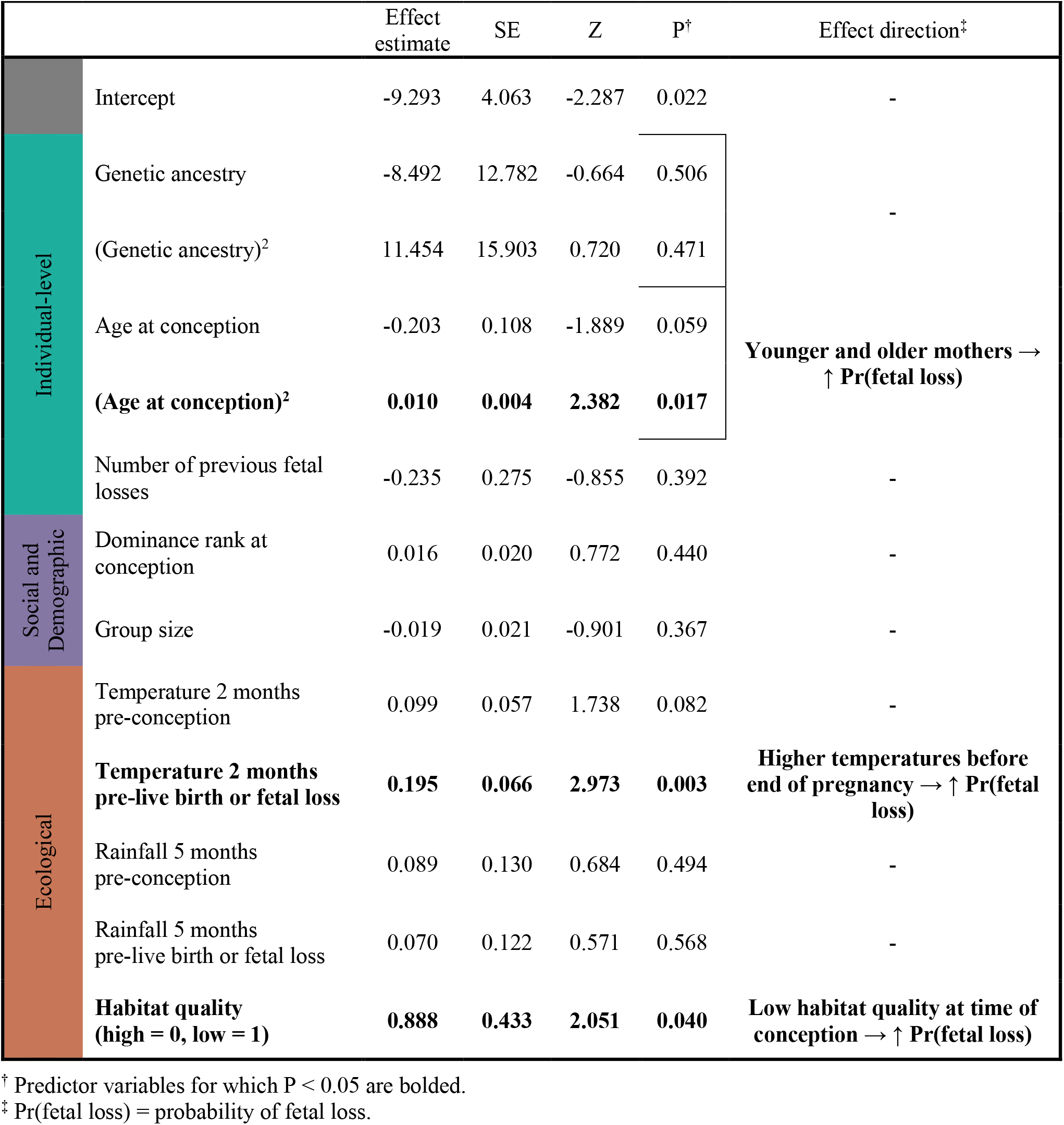
Results from the main logistic regression model predicting fetal loss.

**Figure 3.**
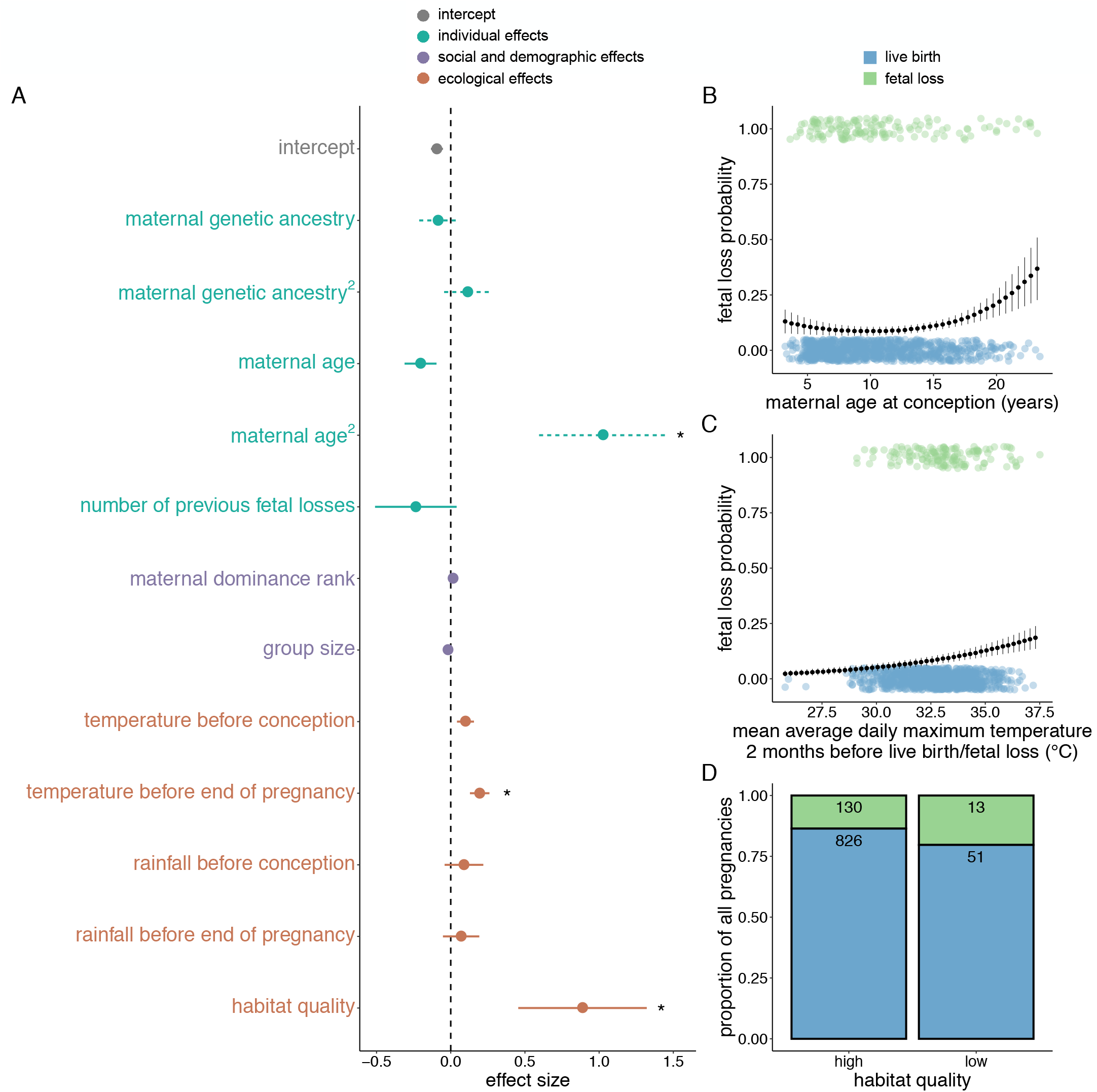
Predictors of fetal loss in wild baboons. **(A)** Effect estimates for the intercept and fixed effects in the main logistic regression model predicting fetal loss. Dots correspond to effect sizes and horizontal lines correspond to effect sizes ± 1 standard error. Dashed horizontal lines correspond to the intercept and genetic ancestry-related predictors (which were divided by 100 to place them on a similar scale as the other predictors, to facilitate visualization) and maternal age^2^ (multiplied by 100 for scaling, for the same reason). Solid horizontal lines correspond to unscaled effect sizes and standard errors. Asterisks denote predictor variables for which P < 0.05. **(B)** The probability of fetal loss as a function of a female’s age at conception. Black dots and vertical lines correspond to the predicted relationship (± 1 standard error for each female age) based on model estimates, assuming average values for all other covariates. Colored dots show fetal loss (y=1; green) or live birth (y=0; blue) for all 1,020 pregnancies (dots are jittered vertically for visibility). **(C)** As in (B), with the probability of fetal loss as a function of temperature two months prior to the end of pregnancy. **(D)** The proportion of all pregnancies that were fetal losses (colored green) or live births (colored blue) as a function of habitat quality. Counts per pregnancy outcome for each type of habitat quality are shown as numbers within each bar.

### Maternal age predicts fetal loss

Our main model identified female age as the only individual-level characteristic that predicted fetal loss (Table 1, Fig. 3A). Younger and older pregnant females were more likely to experience fetal loss than middle aged females (quadratic effect of age: β = 0.010, P = 0.017; Table 1, Fig. 3B). Based on model predictions, females at the oldest age included in our data set (23.2 years) are the most likely to suffer fetal loss (38.6% probability of fetal loss vs. 13.9% for females at the youngest age; Fig. 3B), while females at approximately the median age in the data set (9.73 vs. median 9.49 years) are the least likely to suffer fetal loss (9.3%).

### Ecological factors, but not social status or group size, contribute to variance in pregnancy outcomes

Beyond age effects, which are common in humans and other animals, the primary predictors we identified for fetal loss were related to heat stress and overall habitat quality. Increasing mean maximum temperatures prior to the end of pregnancy predicted an increased chance of fetal loss (β = 0.195, P = 0.003, Table 1, Fig. 3A and 3C; see Beehner, Onderdonk, et al. (2006) for a similar result for first-trimester losses). Elevated mean maximum temperatures prior to conception may also weakly contribute to the probability of fetal loss, although this effect was not statistically significant (β = 0.099, P = 0.082, Table 1, Fig. 3A; see also Tables S1, S3, S5, and S6 for alternative model specifications with similar results). Additionally, pregnancies when females lived in low habitat quality were more likely to end in fetal loss relative to those occurring in high habitat quality (β = 0.888, P = 0.040; Table 1, Fig. 3D), although this effect was not statistically significant under several alternative model specifications (β = 0.857–1.039, P = 0.053–0.103, Table S1, S5, S6). In contrast, social rank, group size, and rainfall levels pre-conception and before the end of pregnancy did not predict fetal loss rates in any version of our models (all P > 0.235; Table 1, Fig. 3A, Tables S1, S3, S5, S6).

## Discussion

Together, our analysis combines long-term data on behavior, demography, and ecology with the ability to track fetal loss to produce the largest study of fetal loss rates in wild primates to date. We found no support for the hypothesis that fetal loss acts to counter free gene flow between yellow baboons and anubis baboons in Amboseli. Instead, our findings show that a female’s genetic ancestry does not predict fetal loss, despite substantial variation in pregnancy outcomes overall. Thus, this aspect of female fertility is unlikely to play a substantial role in maintaining species boundaries in baboons, unless the effects of maternal genetic ancestry are concentrated during the first few weeks of gestation, when we are unable to accurately detect pregnancy. In our data set, the earliest fetal losses occurred at approximately 3 weeks gestational age, which roughly equates to the first 5 weeks of pregnancy in humans. In human women, losses during this time are thought to be associated with spontaneous chromosomal abnormalities, primarily aneuploidies, in the embryo (Larsen, Christiansen, Kolte, & Macklon, 2013; Pinar et al., 2018; but see Brosens et al., 2022). Hybridization can increase aneuploidy rates in some animal systems (Dion-Côté, Symonová, Ráb, & Bernatchez, 2015; Fujiwara, Abe, Yamaha, Yamazaki, & Yoshida, 1997; Hauffe, Giménez, & Searle, 2012; Hu, Huang, Mao, Wang, & Bao, 2013; Sakai et al., 2007). However, hybridization-driven chromosomal abnormalities are not an obvious expectation in baboons, where chromosome numbers are identical in all extant taxa and synteny is thought to be very high (Stanyon et al., 2008).

Our results indicate that the ability of hybrid females to carry a pregnancy to term does not contribute to reproductive isolation between baboon species. However, they do not rule out the possibility that other aspects of conception or pregnancy contribute to reproductive isolation. For instance, we were unable to investigate effects of paternal ancestry in this study because, in a multiply mating species like baboons, paternal identity cannot be confirmed without genetic data, which are not available for miscarried or stillborn fetuses. Thus, it remains possible that genetic ancestry affects fetal loss as a function of the father’s genetic ancestry. This possibility would be consistent with Haldane’s rule, which posits that when species interbreed, hybrids of the heterogametic sex (e.g., fathers) are more likely to suffer fitness costs relative to hybrids of the homogametic sex (e.g., mothers) (Haldane, 1922). Additionally, the interaction between maternal and paternal genetic ancestries could play a role if ancestry combinations at specific regions of the genome negatively interact in the developing fetus (i.e., Bateson-Dobzhansky-Muller incompatibilities, or BDMIs: Bateson, 1909; Dobzhansky, 1936; Muller, 1942). The locations of putative BDMIs can be inferred via genetic scans for local ancestry combinations that are underrepresented in hybrid populations (Payseur & Hoekstra, 2005; Pool, 2015; Schumer et al., 2014), suggesting a potential path forward for testing this hypothesis in hybrid populations where fetal loss and genetic ancestry data are available.

While we found no evidence for genetic ancestry effects, our results do highlight several other sources of variance in pregnancy outcomes in this population. Female age predicted fetal loss, such that the youngest and oldest females experienced the highest rates of fetal loss. In a previous analysis in this population, Beehner, Onderdonk, et al. (2006) found no linear effect of age but did not examine a quadratic effect of age, which may explain the difference between this analysis and the earlier one. However, our results are similar to those described in captive baboons (Schlabritz-Loutsevitch et al., 2008). Indeed, even though captive baboons have much longer life expectancies (Bronikowski et al., 2002), fetal loss rates increase exponentially at ∼14-15 years of age in both Amboseli and in the breeding colony at the Southwest National Primate Research Center (Schlabritz-Loutsevitch et al., 2008), suggesting that the onset of reproductive senescence in female baboons may be relatively insensitive to environmental differences.

Ecological stressors also predicted fetal loss: females living in low habitat quality and exposed to heat stress during pregnancy experienced elevated fetal loss rates. Here again, our findings expand on the work of Beehner, Onderdonk, et al. (2006), who also identified a possible effect of heat stress on fetal loss rates. However, this relationship did not reach statistical significance (P=0.07 in Beehner, Onderdonk, et al. (2006)) and was detectable only in first-trimester pregnancies. Further, their results could have been affected by the systematically elevated maximum daily temperature records during five years of the long-term study (1992-1997; see Methods). Our results, which include a substantially larger sample size (n=1,020 pregnancies versus 656 in Beehner, Onderdonk, et al. (2006)) and corrected temperature data (see Methods), therefore confirm the importance of heat stress for pregnancy outcomes across gestation. Our results also dovetail with evidence from a wide variety of animal taxa on the relationship between heat stress and compromised fertility, starting from germ cell generation through gestation (reviewed in Boni, 2019; Hansen, 2009; Walsh et al., 2019).

In contrast to Beehner, Onderdonk, et al. (2006), which identified an effect of severe drought on pregnancy outcomes, we did not identify an effect of rainfall on pregnancy outcomes. However, our finding that overall habitat quality (not included in Beehner, Onderdonk, et al. (2006)) predicts fetal loss may capture a similar mechanism: a degraded resource base with inadequate food to support successful female reproduction. Notably, exposure to heat stress and drought conditions are expected to become more common in many animals due to accelerating climate change (Fuller et al., 2021; Walsh et al., 2019). Our results suggest that these changes may not only compromise habitat quality, but also alter vital rates for primate populations to decrease mean population fitness.

Our findings also reinforce the value of baboons as models for human reproduction (Bauer, 2015; D’Hooghe, Kyama, & Mwenda, 2009; Hendrickx & Peterson, 2009; Honoré & Tardif, 2009; Nathanielsz et al., 2009). Specifically, in addition to sharing slow life histories, little to no reproductive seasonality, and similarities in reproductive biology (e.g., size and anatomy of the internal reproductive tract, morphology of the placenta, incidence of gynecological diseases; reviewed in VandeBerg, Williams-Blangero, & Tardif, 2009), our study suggests that fetal loss in baboons exhibits parallels to fetal loss in humans (Schlabritz-Loutsevitch et al., 2008). The rate of fetal loss in Amboseli—approximately 1.4 out of ten pregnancies (Beehner, Onderdonk, et al., 2006 and this study)—is similar to some estimates of miscarriage rates for clinically recognized pregnancies in humans (e.g., ∼10-20% after implantation: Dimitriadis et al., 2020; Pinar et al., 2018). Moreover, abiotic environmental stressors, including high temperatures during pregnancy, have recently gained attention in potentially explaining adverse birth outcomes in humans, such as fetal loss (e.g., Hajdu & Hajdu, 2021; Kanner et al., 2020; Strand, Barnett, & Tong, 2012; Syed, O’Sullivan, & Phillips, 2022; but see Asamoah, Kjellstrom, & Ostergren, 2018). For example, in low-risk pregnant women in Utah, extreme heat exposure (>90^th^ temperature percentile) increased the odds of stillbirth by ∼5-fold compared to exposure to moderate temperatures (Kanner et al., 2020). Further, age-related miscarriage risk in human women is thought to also follow a “J-shaped curve,” with higher rates in very young women and increasing rates as women age (Brosens et al., 2022). Notably, in humans, age-related patterns of miscarriage also suffer from sociocultural biases that influence both pregnancy outcomes and the age of first pregnancy (Cohen, 2014; Santelli, Song, Garbers, Sharma, & Viner, 2017). Our novel observation that a similar “J-shaped curve” occurs in wild baboons therefore provides preliminary evidence that the pattern observed in humans may be in part due to evolutionarily conserved reproductive biology. Importantly, in humans, the causes of a large fraction of pregnancy failures remain unclear despite decades of biomedical research (Pinar et al., 2018). Thus, studies in wild non-human primates may therefore present valuable systems for understanding biological drivers of adverse pregnancy outcomes in humans.

## Acknowledgements

This research was supported by the U.S. National Science Foundation (NSF) and the U.S. National Institutes of Health (NIH), currently through NSF IOS 1456832 (S.C.A), NSF BCS-2018897 (J.T., A.S.F.), NIH R01AG053308 (S.C.A), R01AG053330 (E.A.A.), R01AG071684 (E.A.A.), R01HD088558 (J.T.), R01AG075914 (J.T.), and P01AG031719 (S.C.A.). A.S.F. was supported by NSF GRFP (DGE #1644868) and NIH T32GM007754. We are grateful to Duke University, the University of Notre Dame, and Princeton University for financial and logistical support. In Kenya, our research was approved by the Wildlife Research Training Institute (WRTI), Kenya Wildlife Service (KWS), the National Commission for Science, Technology, and Innovation (NACOSTI), and the National Environment Management Authority (NEMA). We also thank the University of Nairobi, Institute of Primate Research (IPR), the National Museums of Kenya, the Amboseli-Longido pastoralist communities, the Enduimet Wildlife Management Area, Ker & Downey Safaris, Air Kenya, and Safarilink for their assistance in Kenya. Particular thanks go to Jeanne Altmann for her fundamental contributions to research on the Amboseli baboons, the Amboseli Baboon Research Project field team (R. S. Mututua, S. Sayialel, J. K. Warutere, I. L. Siodi, G. Marinka, B. Oyath) for their data collection in the field, the Amboseli Baboon Research Project camp staff for their support during fieldwork, and T. Wango and V. Oudu for their assistance in Nairobi. Finally, we thank K. Pinc, N. H. Learn, and J. B. Gordon for managing the Amboseli Baboon Research Project long-term database. For a complete set of acknowledgments of funding sources, logistical assistance, and data collection and management, please visit http://amboselibaboons.nd.edu/acknowledgements/. Any opinions, findings, and conclusions expressed in this material are those of the authors and do not necessarily reflect the views of our funding bodies.

## Author Contributions

Conceptualization, E.A.A., J.T., S.C.A.; Methodology, A.S.F., E.A.A., J.T., P.O.O.; Investigation, A.S.F., E.A.A., J.T., P.O.O.; Formal Analysis, A.S.F. and P.O.O.; Resources, A.W.N., C.N.K., E.A.A., J.T., S.C.A.; Data Curation, E.A.A., J.T., S.C.A.; Writing – Original Draft, A.S.F. and J.T.; Writing – Reviewing & Editing, A.S.F., A.W.N., C.N.K., E.A.A., J.T., P.O.O., S.C.A.; Visualization, A.S.F.; Supervision, A.W.N., C.N.K., E.A.A., J.T., S.C.A.; Project Administration, A.W.N., C.N.K., E.A.A., J.T., S.C.A.; Funding Acquisition, A.S.F., E.A.A., J.T., S.C.A.

## Supplementary Information

**Figure S1.**
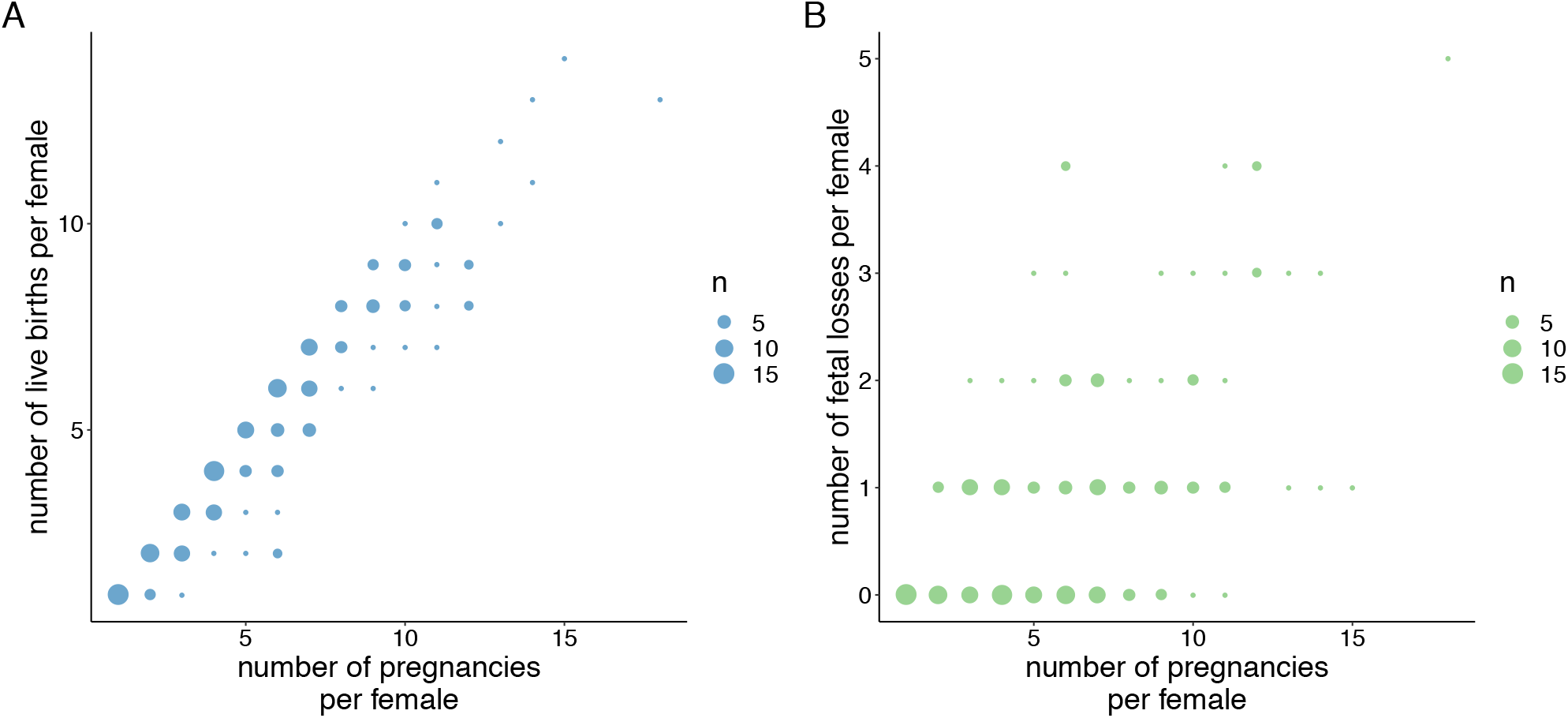
The total number of pregnancies per female predicts **(A)** the total number of live births per female (β = 0.824, p < 10^−89^) and **(B)** the total number of fetal losses per female (β = 0.176, p < 10^−14^). Each value corresponds to a unique female in the data set (e.g., n=5 is 5 unique females).

**Table S1.**
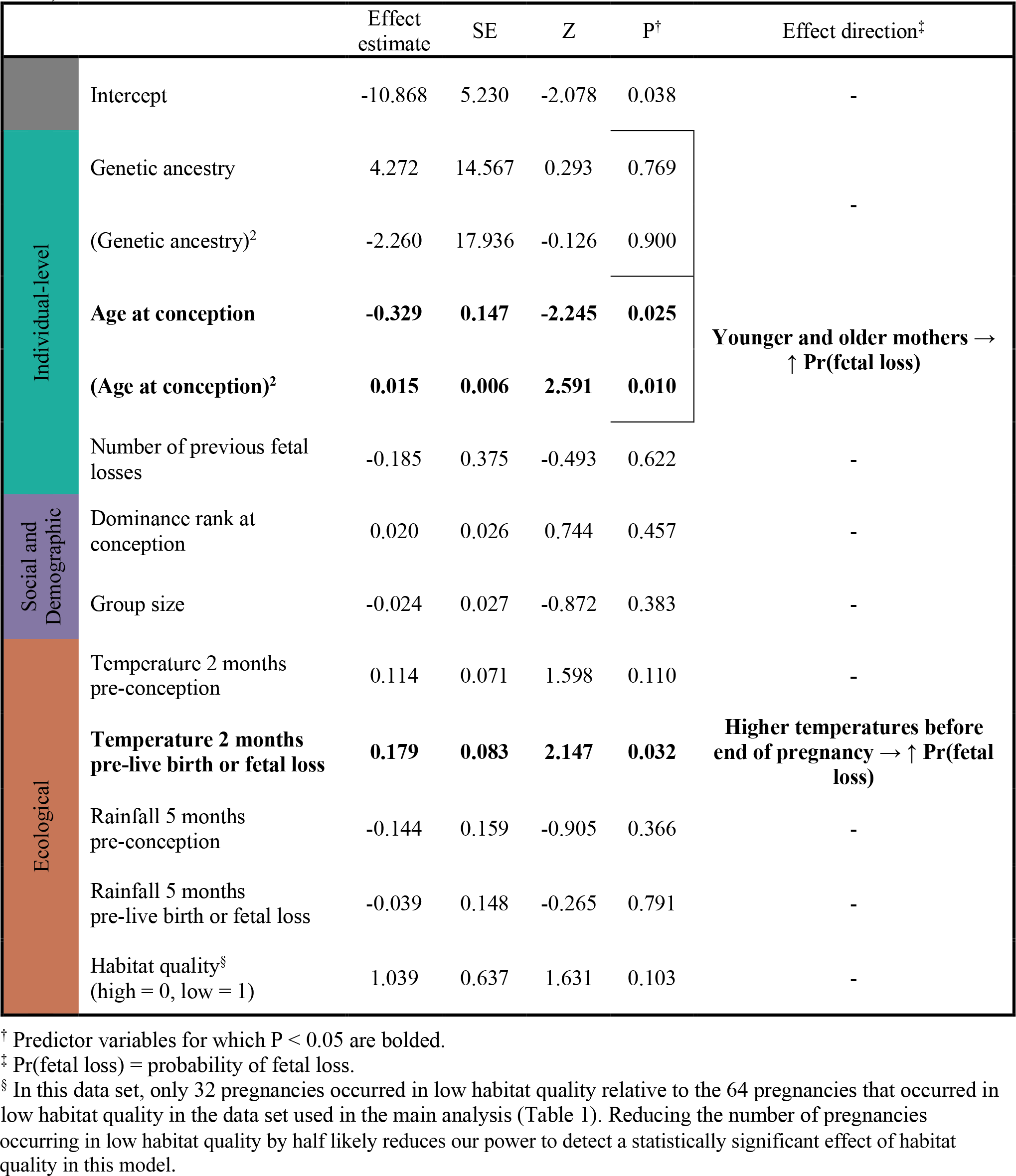
Results from a logistic regression model predicting fetal loss using the most conservative inclusion criteria. We excluded females in non-wild feeding social groups, females with birthdates estimated with greater than a few days’ error, pregnancies that overlapped the 2009 hydrological year (a severe drought), pregnancies that occurred during several periods of reduced data collection, pregnancies that occurred during social group fissions and fusions, and pregnancies with conception and end dates that were not known to within a few days’ error (n=676 pregnancies, 142 females). Results are qualitatively similar to those in Table 1 (excluding an effect of habitat quality that is statistically significant in Table 1; see note § below table).

**Table S2.** Pearson’s product-moment correlation (*r*) between continuous predictor variables in the data set and p-values in parentheses.

|  | Age at conception | Number of previous fetal losses | Dominance rank at conception | Group size | Temperature pre-conception | Temperature pre-live birth or fetal loss | Rainfall pre-conception | Rainfall pre-live birth or fetal loss |
| --- | --- | --- | --- | --- | --- | --- | --- | --- |
| Genetic ancestry | 0.000<br>(0.988) | 0.046<br>(0.144) | 0.018<br>(0.556) | 0.114<br>**(<0.001)** | -0.062<br>(0.046) | -0.026<br>(0.411) | -0.007<br>(0.825) | 0.018<br>(0.567) |
| Age at conception |  | 0.422<br>**(<0.001)** | -0.011<br>(0.727) | 0.019<br>(0.554) | 0.003<br>(0.928) | 0.004<br>(0.907) | -0.026<br>(0.416) | 0.033<br>(0.291) |
| Number of previous fetal losses |  |  | 0.006<br>(0.844) | -0.006<br>(0.854) | 0.001<br>(0.962) | 0.022<br>(0.490) | -0.023<br>(0.456) | 0.035<br>(0.268) |
| Dominance rank at conception |  |  |  | 0.468<br>**(<0.001)** | 0.048<br>(0.124) | -0.041<br>(0.191) | 0.005<br>(0.872) | -0.029<br>(0.352) |
| Group size |  |  |  |  | 0.011<br>(0.719) | -0.041<br>(0.194) | 0.040<br>(0.204) | -0.039<br>(0.216) |
| Temperature pre-conception |  |  |  |  |  | -0.157<br>**(<0.001)** | 0.064<br>(0.041) | -0.204<br>**(<0.001)** |
| Temperature pre-live birth or fetal loss |  |  |  |  |  |  | -0.325<br>**(<0.001)** | 0.018<br>(0.563) |
| Rainfall pre-conception |  |  |  |  |  |  |  | -0.385<br>**(<0.001)** |

P-values < 0.01 are bolded and P < 0.05 are italicized.

**Table S3.**
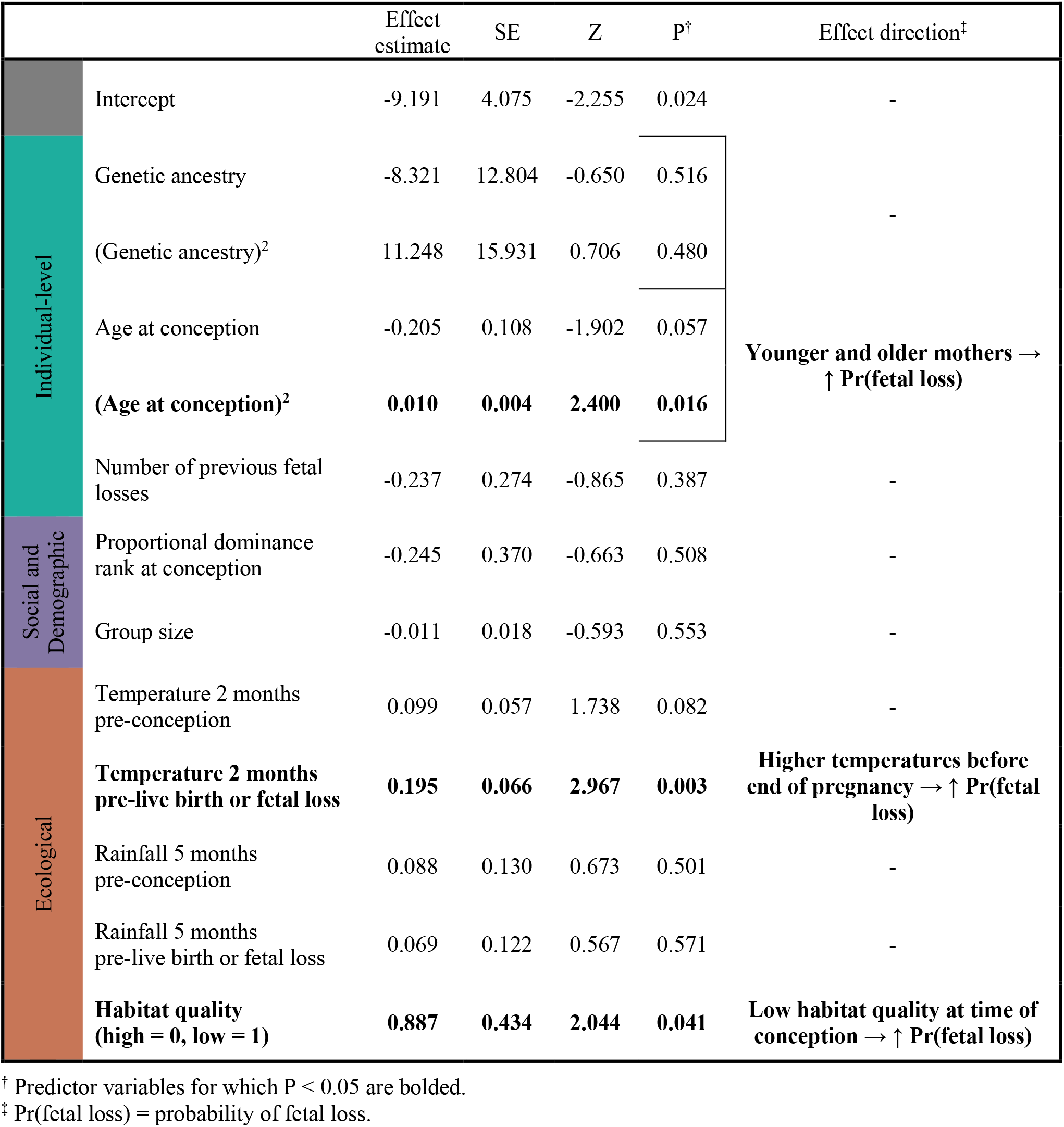
Results from a logistic regression model predicting fetal loss, using a mother’s proportional rank instead of ordinal rank at the time of conception as a predictor variable. Results are qualitatively similar to results presented in Table 1.

**Table S4.** Data on pregnancy outcomes and all predictor variables tested in our models. Available in Supplementary Information excel file.

**Table S5.**
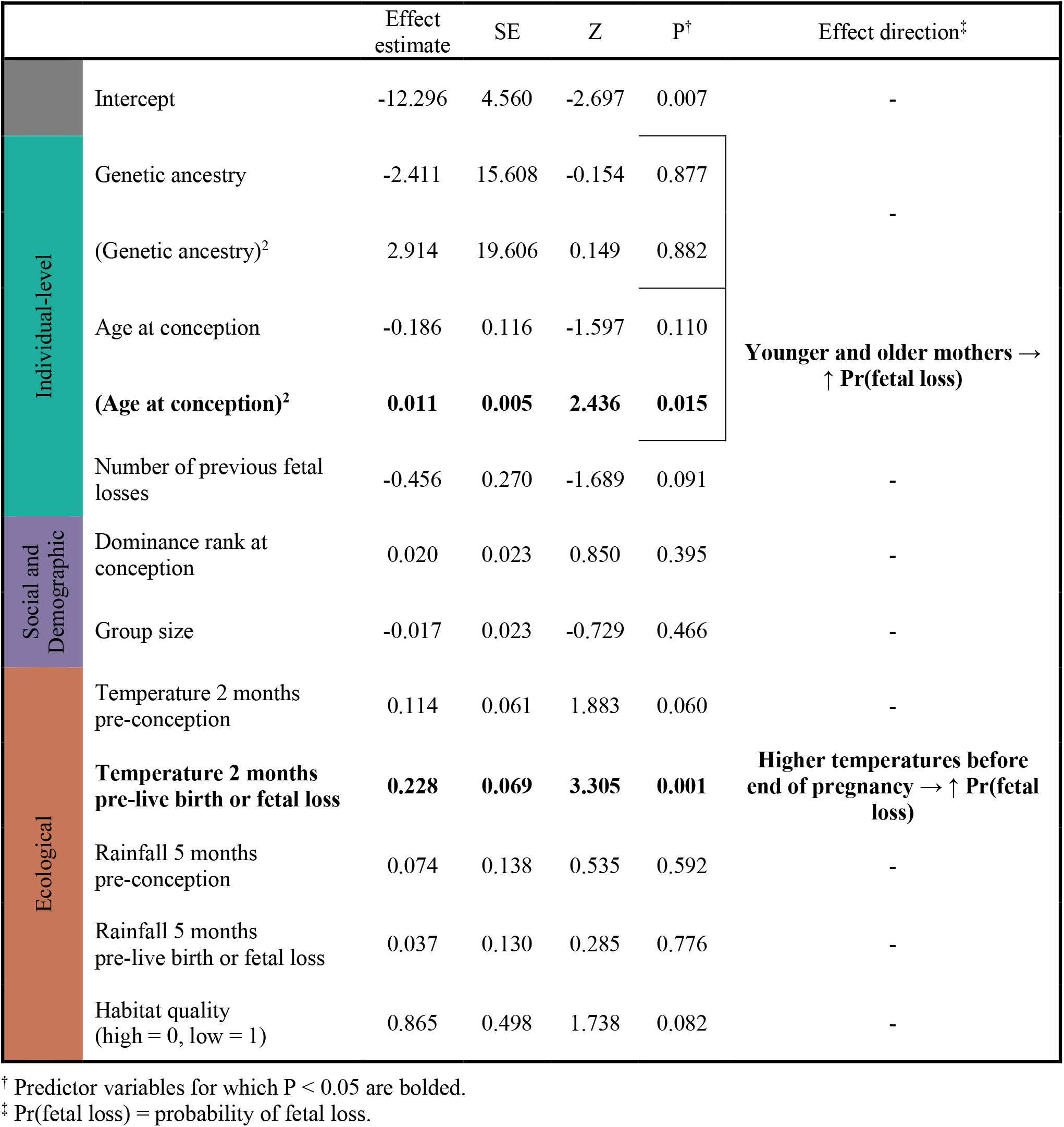
Results from a logistic regression model predicting fetal loss excluding fetal losses due to feticide or presumed feticide (n=8). Results are qualitatively similar to results presented in Table 1 (excluding an effect of habitat quality that is statistically significant in Table 1).

**Table S6.**
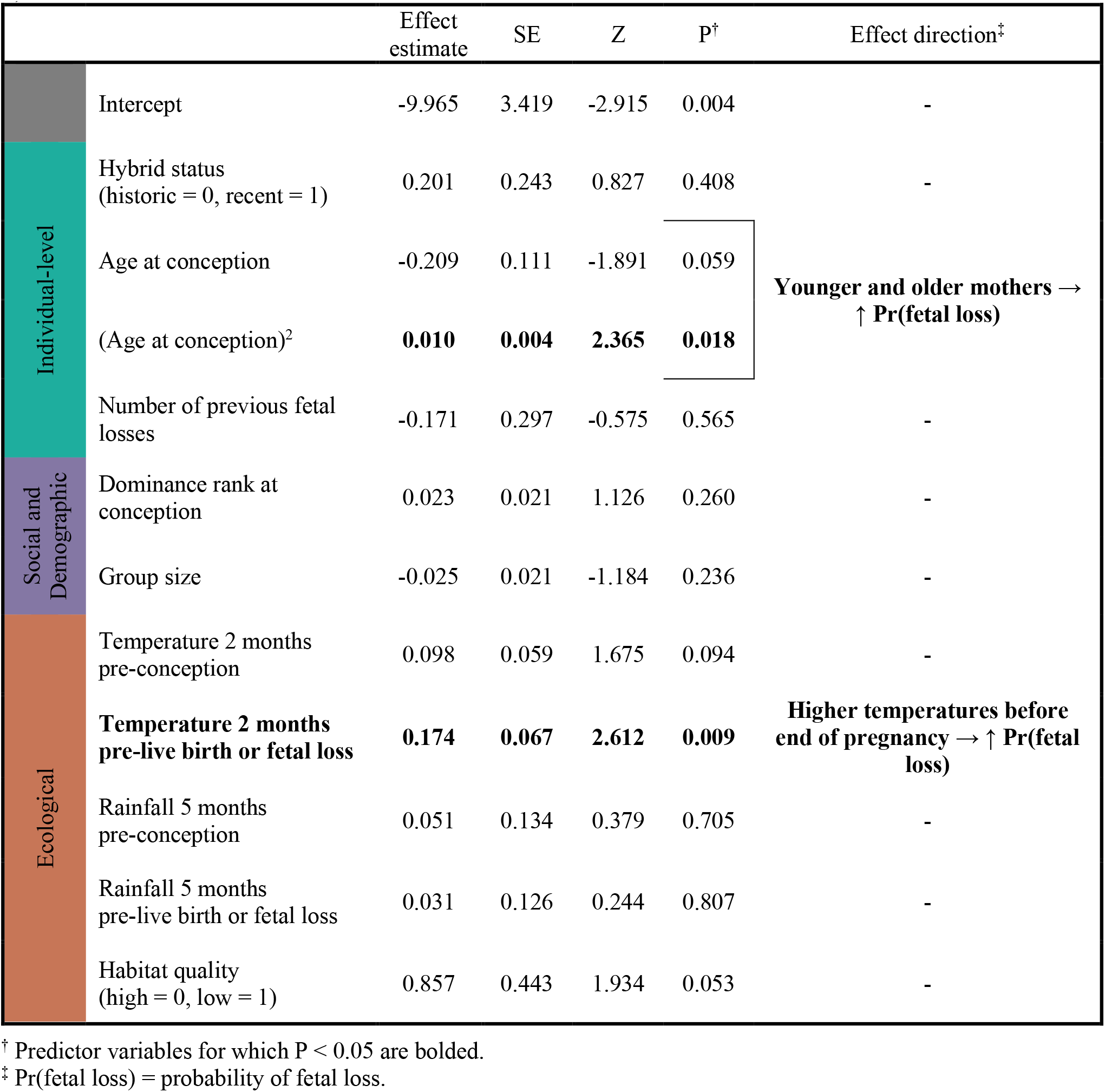
Results from a logistic regression model predicting fetal loss substituting hybrid status for genome-wide ancestry as a predictor variable. Results are qualitatively similar to results presented in Table 1 (excluding an effect of habitat quality that is statistically significant in Table 1).

